# Overexpression of Meis factors in late-stage retinal progenitors yields complex effects on temporal patterning and neurogenesis

**DOI:** 10.1101/2025.05.01.651492

**Authors:** Patrick Leavey, Lizhi Jiang, Nicole Pannullo, Clayton Santiago, Seth Blackshaw

## Abstract

The vertebrate retina serves as a model for studying neurogenesis and cell fate specification, with retinal progenitor cells following a tightly regulated temporal sequence to generate distinct cell types. Meis1 and Meis2 are transcription factors implicated in early retinal development, but their role in late-stage RPCs remains poorly understood. Here, we investigate whether *Meis1* and *Meis2* overexpression in postnatal mouse RPCs can alter temporal identity and induce early-born cell types. Using electroporation and single-cell RNA sequencing, we find that while these factors modestly upregulate early-stage gene regulatory network components, they do not repress late-stage transcription factors or induce early-born retinal cells. *Meis1* overexpression reduces proliferation and inhibits neurogenesis, whereas *Meis2* overexpression accelerates neurogenic progression without altering fate commitment. Our findings suggest that overexpression of *Meis1* and *Meis2* modulate largely non-overlapping aspects of temporal identity and neurogenic competence but are insufficient to fully reprogram late-stage progenitors. These results have implications for regenerative strategies aimed at reprogramming retinal cells for therapeutic purposes.

## Introduction

The vertebrate retina, as a lateral extension of the embryonic neural tube, provides a simplified yet integral model for studying neurogenesis and cell fate specification in the central nervous system (CNS) (X. Zhang et al. 2023). It consists of six major neuronal and one glial cell type, all of which arise from retinal progenitor cells (RPCs) in a temporally regulated sequence. Birthdating studies have shown that retinal ganglion cells are generated first, followed by horizontal cells, GABAergic amacrine cells, cone photoreceptors, non-GABAergic amacrine cells, rod photoreceptors, bipolar cells, and finally Müller glia (Cepko 2014; Maurange 2020). Clonal analyses of isolated RPCs demonstrate that these changes in developmental potential are largely intrinsic (Gomes et al., 2011; Cayouette et al., 2003), driven by dynamic transcription factor expression that governs temporal competence (Gomes et al. 2011; Cayouette, Barres, and Raff 2003). While several key transcription factors, including *Ikzf1/4, Nfia/b/x, Foxp1*, and *Caszl*, have been identified as regulators of temporal identity (Boudreau-Pinsonneault et al. 2023; Clark et al. 2019; J. Zhang et al. 2023; Mattar et al. 2015), much remains unknown about how these factors coordinate developmental transitions.

Recent single-cell RNA sequencing (scRNA-Seq) and chromatin accessibility (scATAC-Seq) studies have identified additional candidate regulators of temporal patterning in the developing retina (Clark et al. 2019; Lyu et al. 2021; Lu et al. 2020). Early-stage TFs exhibit high activity in retinal neuroepithelial cells prior to neurogenesis and gradually decline as early-born cell types, such as ganglion cells and cone photoreceptors, are generated. In contrast, the late-stage GRN becomes increasingly active from embryonic day (E) 17 in mice (gestational weeks 11–13 in humans), persisting into postmitotic Müller glia, the final retinal cell type to emerge (X. Zhang et al. 2023; Lyu et al. 2021). These GRNs are composed of interdependent TFs that reinforce their own expression while repressing factors from the opposing network, yet the functional roles of many predicted regulators remain unexplored., most of which have not been systematically studied for their ability to regulate temporal patterning.

*Meis1* and *Meis2* are transcription factors predicted to contribute prominently to the early-stage GRN (Lyu et al. 2021). Both genes are selectively expressed in early-stage RPCs and directly regulate transcription of multiple early-stage transcription factors. Genetic studies have demonstrated that Meis factors are essential for many different aspects of early retinal development. In zebrafish, *meis1* is essential for both dorso-ventral and naso-temporal patterning of the retina prior to the onset of neurogenesis (Erickson, French, and Waskiewicz 2010), while *Meis1* deficiency in mice disrupts neuroretinal formation and delays the onset of neurogenesis (Marcos et al. 2015; Heine et al. 2008). Combined loss of *Meis1/2* completely disrupts neuroepithelial proliferation and neurogenesis, arresting progenitors at an undifferentiated state (Dupacova et al. 2021). The functions of *Meis1/2* during later stages of neurogenesis have been less extensively investigated. Loss of function of *Meis1* following the onset of retinal neurogenesis inhibits expression of *Foxn4*, which is necessary for generation of early-born horizontal and amacrine cells (Islam et al. 2013). *Meis2* is transiently expressed in developing cone photoreceptors and non-GABAergic amacrine cells and persists in a subset of mature GABAergic amacrine cells (Bumsted-O’Brien et al. 2007). However, whether ectopic *Meis1/2* expression in late-stage RPCs can reprogram temporal identity and induce early-born cell types remains unknown. This question is particularly relevant given recent efforts to induce neurogenesis from mature Müller glia following overexpression of neurogenic bHLH factors or Oct4, inhibition of Notch signaling, or loss of function of the late-stage GRN components *Nfia/b/x* (Todd et al. 2022; Wohlschlegel et al. 2023; Hoang et al. 2020; Le, Vu, et al. 2024; Le, Awad, et al. 2024). While these strategies have successfully converted Müller glia into interneurons, they have largely failed to produce photoreceptors.

In this study, we use electroporation and scRNA-Seq to investigate whether *Meis1, Meis2*, or their combined overexpression can induce postnatal RPCs to adopt an early-stage temporal identity and generate early-born cell types. While overexpression of these factors modestly increased expression of early-stage GRN components, it did not suppress late-stage TFs. *Meis1* overexpression reduced proliferation and inhibited neurogenesis, whereas *Meis2* accelerated neurogenic progression. However, neither factor induced ectopic formation of early-born retinal cells. These findings suggest that *Meis1/2* can modulate aspects of temporal identity and neurogenic competence but are insufficient to fully reprogram late-stage progenitors or drive robust neurogenesis.

## Results

### *Meis1* and *Meis2* expression in retinal progenitor cells

To assess the temporal dynamics of *Meis1* and *Meis2* expression in primary and neurogenic retinal progenitor cells (RPCs), we analyzed previously published single-cell RNA sequencing (scRNA-Seq) datasets from developing mouse and human retina (Clark et al. 2019; Zuo et al. 2024) (Fig. 1a,b; S1). Consistent with prior studies (Heine et al. 2008), both genes were highly expressed in early-stage primary RPCs, peaking at embryonic day (E) 12 in mice, coinciding with the onset of retinal neurogenesis (Riesenberg et al. 2009). A similar pattern was observed in fate-restricted neurogenic RPCs, with expression also peaking at E12. In the human retina, *MEIS1* levels declined progressively from gestational week (GW) 8 onward, whereas *MEIS2* exhibited an opposing trend, peaking at GW17 (Fig. S2a,b; S3a,b).

**Figure 1:**
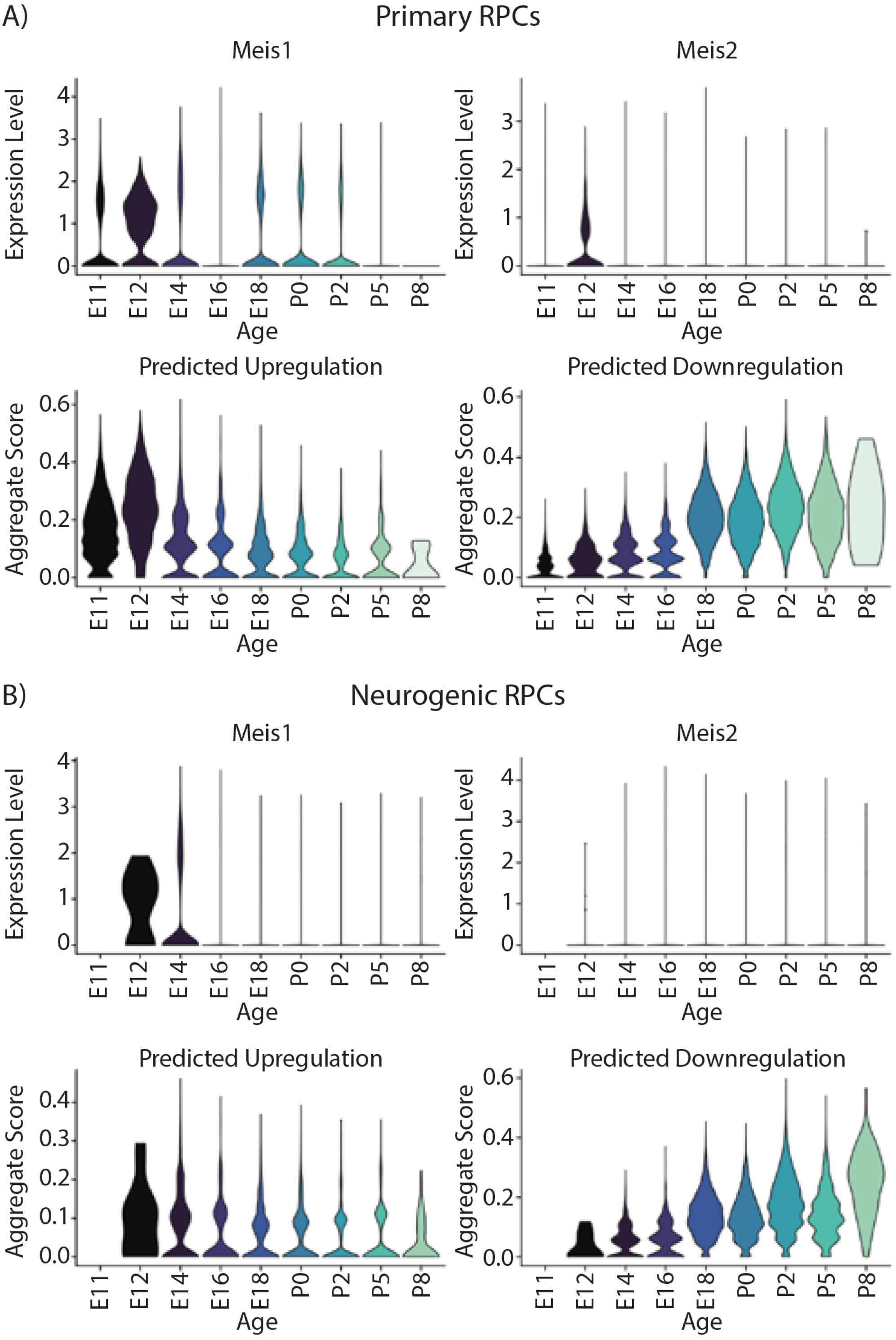
Expression of *Meis1/2* and expression of genes predicted to be directly upregulated or downregulated by Meis factors in mouse retina. A) Primary retinal progenitor cells and B) neurogenic retinal progenitors were analyzed from the dataset published in Clark et al 2019. Violin expression plots for Meis1 and Meis2. Violin scoring plots for predicted upregulated and downregulated GRNs from Lyu et al 2021.

Previously, our group identified evolutionarily conserved gene regulatory networks (GRNs) controlling temporal patterning in mouse and human retinal progenitors, including predicted targets of *Meis1* and *Meis2* (Lyu et al. 2021). In mice, genes activated by these factors were enriched in early-stage progenitors, whereas genes predicted to be repressed by *Meis1/2* were upregulated at later stages, mirroring the dynamic expression of these TFs (Fig. 1a,b; S1a,b). In the human retina, genes repressed by *MEIS1/2* exhibited similar late-stage upregulation, but activation of target genes in early RPCs was less pronounced (Fig. S2a,b; S3a,b).

### *MEIS1* and *MEIS2* overexpression in late-stage mouse retinal progenitor cells

To investigate whether *MEIS1* and *MEIS2* influence RPC temporal identity and neurogenesis, we electroporated P0 mouse retinas ex vivo with pCAGIG-based plasmids expressing full-length human *MEIS1, MEIS2*, or a 50:50 mixture of both. Control retinas were electroporated with empty pCAGIG vectors (Fig. 2a). Retinas were cultured until P2 or P5, after which GFP-positive cells were isolated via fluorescence-activated cell sorting (FACS) (Fig. S4a,b). scRNA-Seq analysis was performed on duplicate samples for each condition, pooling multiple retinas (n=12–16 per condition). All cells included in the analysis expressed at least one copy of *GFP* mRNA (Fig. S4c–e).

**Figure 2:**
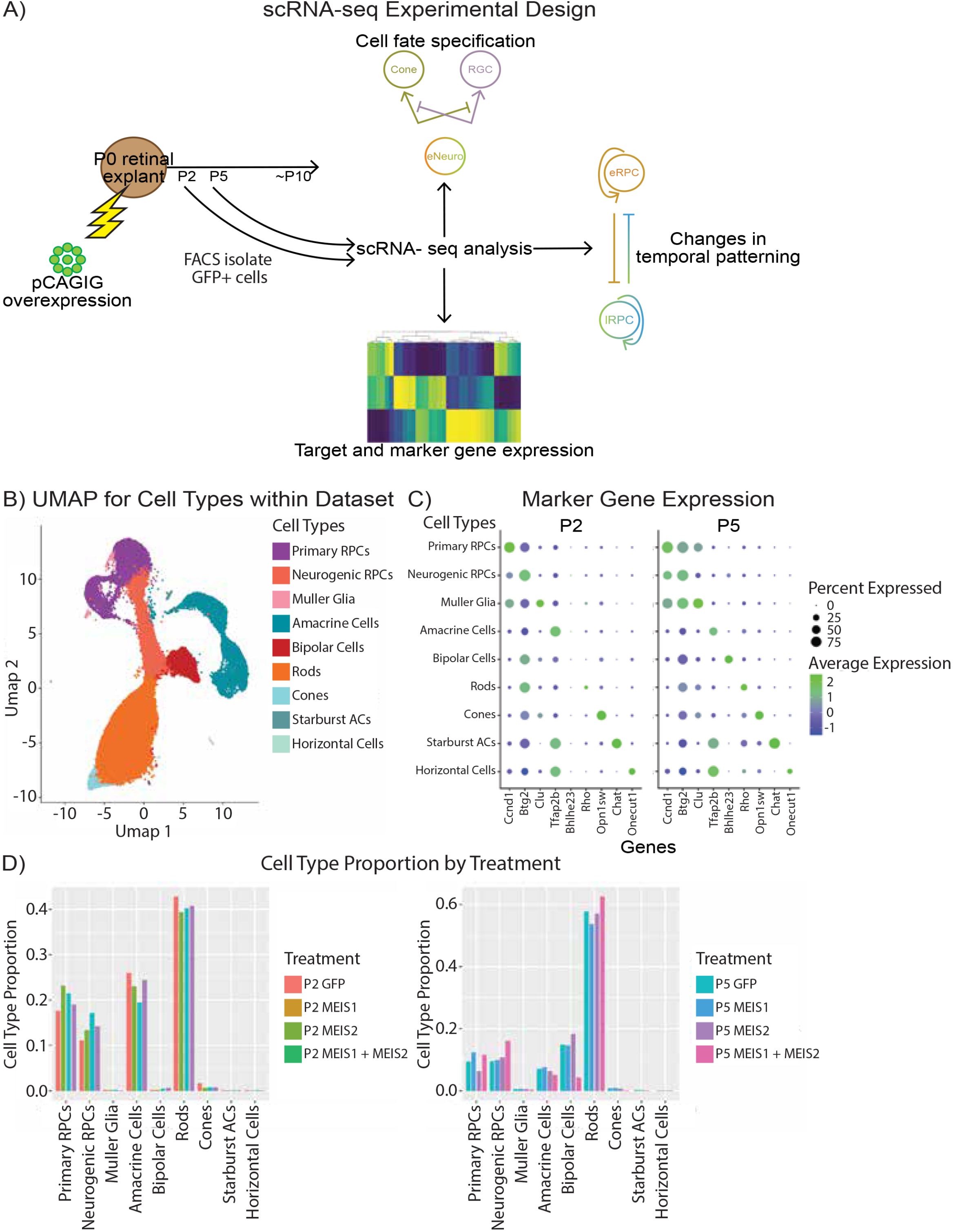
Generating scRNA-Seq datasets for wild type murine retinas overexpressing either control pCAGIG or *MEIS1/2* constructs. A) Schematic for ectopic *MEIS1/2* overexpression and isolation of GFP-positive electroporated cells from P0 retinal explants. B) Uniform manifold approximation (UMAP) and projection for the scRNA-Seq dataset created in A. C) DotPlot for the cell types present at P2 and P5 within the scRNA-Seq dataset and their associated marker genes. D) Proportion of each cell type present in each of the treatment conditions.

Cell types were identified via UMAP clustering and marker gene expression (Fig. 2b,c; Fig. S5, Supplemental Table 1). The composition of electroporated cell populations was consistent with previous studies (Clark et al. 2019; Lyu et al. 2021), with primary and neurogenic RPCs and amacrine cells predominant at P2, while rods and bipolar cells became more abundant at P5 (Fig. 2d). Minimal numbers of Müller glia were detected, and rare GFP-positive cones, starburst amacrine, and horizontal cells were likely contaminants. Small numbers of GFP-negative microglia were also removed before analysis.

*MEIS1/2* overexpression did not alter cell type composition at either P2 or P5 (Fig. 2d), indicating that these factors do not independently induce early-born cell types such as retinal ganglion cells, cones, horizontal cells, or GABAergic amacrine cells. They also neither promoted nor inhibited neurogenesis nor altered the maintenance of primary or neurogenic RPC pools.

However, potential effects on temporal patterning or gene expression required further investigation.

### *MEIS1* and *MEIS2* regulate expression of genes specific to early-stage retinal progenitors

To assess whether *MEIS1* and *MEIS2* regulate their own expression, we examined their auto- and cross-regulation. Overexpression of *MEIS1* activated endogenous *Meis1* but had no effect on *Meis2*, while *MEIS2* induced *Meis2* expression but did not affect *Meis1* (Fig. 3a). A comprehensive analysis of gene expression changes induced by MEIS overexpression (log2 fold change > 0.5, p<0.00001) revealed that both *MEIS1* and *MEIS2* upregulated the early-stage RPC marker *Hmga1* (Fig. 3b,c; Fig. S6a, Supplemental Table 2). *MEIS2* specifically upregulated early-stage factors *Pbx1, E2f1*, and *Foxp4*, as well as the late-stage factor *Nfib*, while *MEIS1* upregulated *Hes1* and *Hes5*, key Notch pathway regulators. *MEIS1* downregulated the neurogenic *bHLH* factors *Neurod1* and *Neurod4*, the latter of which was also downregulated by *MEIS2* (Fig. 3b,c). Notably, co-expression of both factors reversed these effects, increasing *Neurod1* and *Neurod4* expression (Fig. S6a, Supplemental Table 2).

**Figure 3:**
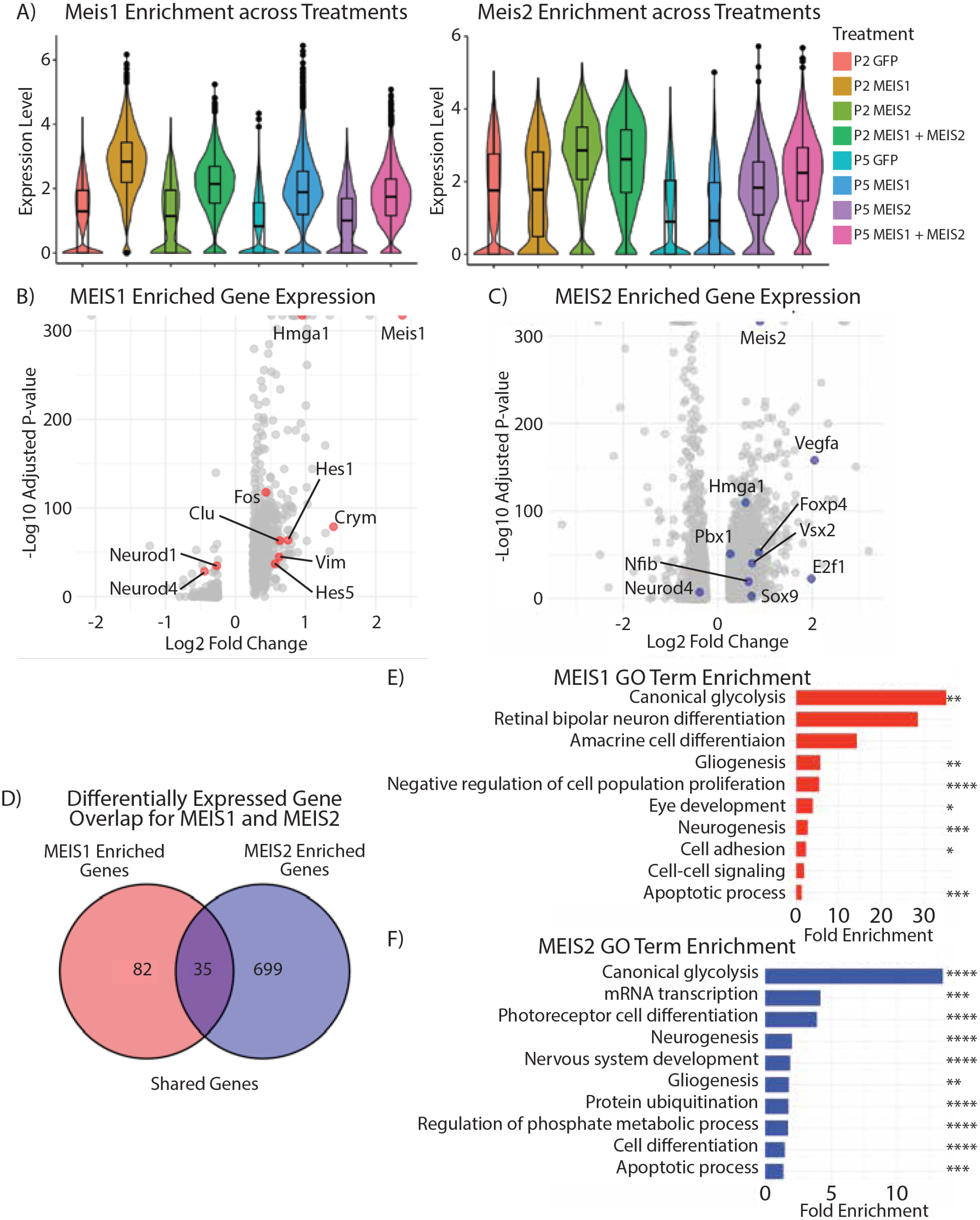
Changes in gene expression induced by *MEIS1/2* overexpression. A) Violin plots showing how each treatment affects the expression of endogenous *Meis1* or *Meis2* across the full dataset. B) Volcano plot generated from differentially-expressed genes between the P2 and P5 control treated samples and the P2 and P5 samples overexpressing either *MEIS1* or C) *MEIS2*. D) Volcano plot generated from the differentially expressed genes between the P2 and P5 control treated samples and the P2 and P5 samples overexpressing *MEIS2*. E) Venn diagram showcasing the overlap in the genes expressed at a 0.5 Log2 Fold change between P2 and P5 *MEIS1* and *MEIS2* overexpressing samples. E) Curated GO Terms selected from generated list from the 117 genes with a 0.5 Log2 Fold increase with *MEIS1* overexpression. F) Curated GO Terms selected from the 734 genes with a 0.5 Log2-fold increase with *MEIS2* overexpression. P-values are indicated by * number. * p< 0.05, ** p< 0.01, *** p< 0.001, and **** p< 0.0001.

Gene ontology (GO) analysis of differentially expressed genes revealed distinct functional categories. *MEIS1* targets were enriched for glycolysis, gliogenesis, neurogenesis, eye development, and apoptosis (Fig. 3e), whereas *MEIS2* targets included glycolysis, gliogenesis, neurogenesis, apoptosis, cell differentiation, and mRNA transcription (Fig. 3f).

Despite substantial functional overlap, the specific genes regulated by each factor showed limited overlap (Fig. S6b).

Finally, we tested whether MEIS factor overexpression led to induction of genes specific to individual postmitotic retinal cell types in either primary or neurogenic RPCs (Fig. S7, S8).

We observed that *MEIS2*, both individually or in combination with *MEIS1*, activated expression of cone-specific genes at both primary and neurogenic RPCs at P2 and P5, and likewise activated amacrine cell-specific genes at P5. No other significant changes in cell-specific gene expression were observed.

### Effects of overexpression of *MEIS1* and *MEIS2* on retinal progenitor cell temporal identity and differentiation

To determine whether *MEIS1* and *MEIS2* influence RPC temporal identity, we calculated a temporal index based on enrichment for early- and late-stage markers (Lyu et al. 2021).

Genes previously predicted to be activated by MEIS factors were upregulated in all conditions at P2, except for *MEIS1* in neurogenic RPCs (Fig. 4a,b; Fig. S9). Surprisingly, neither *MEIS1* nor *MEIS2* repressed predicted targets; instead, these genes exhibited modest but statistically significant upregulation, particularly at P2 (Fig. 4a,b; Fig. S9).

**Figure 4:**
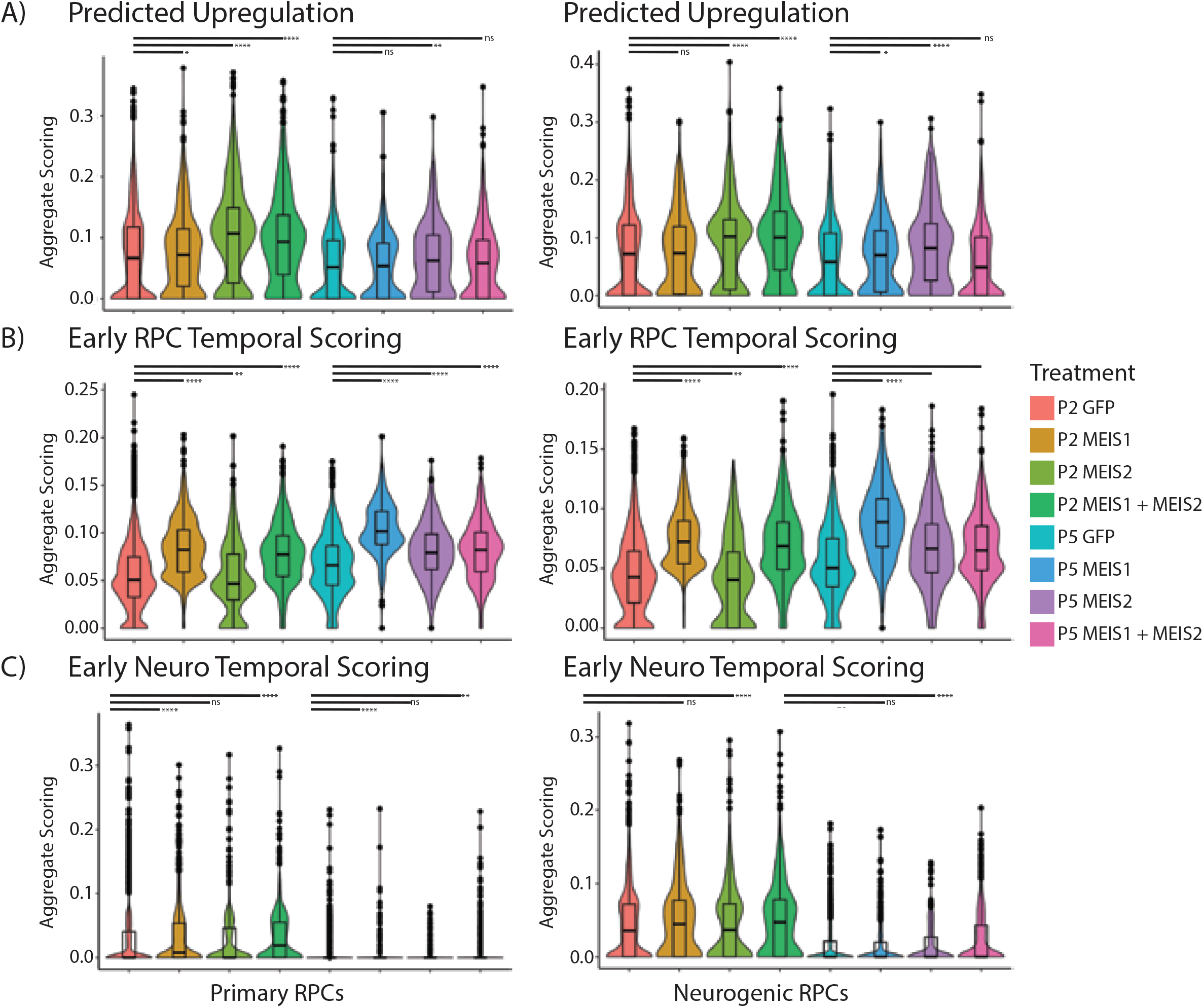
MEIS overexpression’s impact on predicted target genes and progenitor cell temporal identity. A) AUCell R package was used to generate composite scores for genes predicted to be upregulated by *MEIS1* and/or *MEIS2* during retinal development . Split into separate violin plots for primary and neurogenic retinal progenitor cells. B) Scoring violin plots in RPCs for genes expressed in primary RPCs during early developmental stages. C) Scoring violin plots for genes expressed in neurogenic RPCs during early developmental stages. P-values are indicated by * number. ns = not significant, * p< 0.05, ** p< 0.01, *** p< 0.001, and **** p< 0.0001.

Analysis of early- and late-stage marker genes revealed that *MEIS1* strongly induced early-stage markers in both primary and neurogenic RPCs, while *MEIS2* showed this effect only at P5. Co-expression of *MEIS1* and *MEIS2* enhanced early-stage identity across all conditions. However, none of the tested constructs repressed late-stage markers; instead, *MEIS1* and *MEIS1+MEIS2* unexpectedly upregulated these genes (Fig. 4b-c, Fig. S9). In primary RPCs, *MEIS2* but not *MEIS1* repressed Müller glia-specific genes at P2 while inducing amacrine/horizontal and cone markers (Fig. S7). Further analysis showed that *MEIS2* upregulated amacrine/horizontal and cone-specific genes at P2 and P5 in neurogenic RPCs, with additional activation of bipolar-specific genes at both time points (Fig. S8).

To assess whether *MEIS1/2* overexpression altered RPC differentiation timing, we performed pseudotime analysis using AUCell values for early-stage markers. At P2, *MEIS2* accelerated differentiation in the youngest primary RPCs, with both *MEIS1* and *MEIS2* exerting this effect at P5. In neurogenic RPCs, *MEIS2* accelerated differentiation at both P2 and P5, whereas *MEIS1* had no effect. No treatment affected the oldest progenitor populations (Fig. 5; Fig. S8).

**Figure 5:**
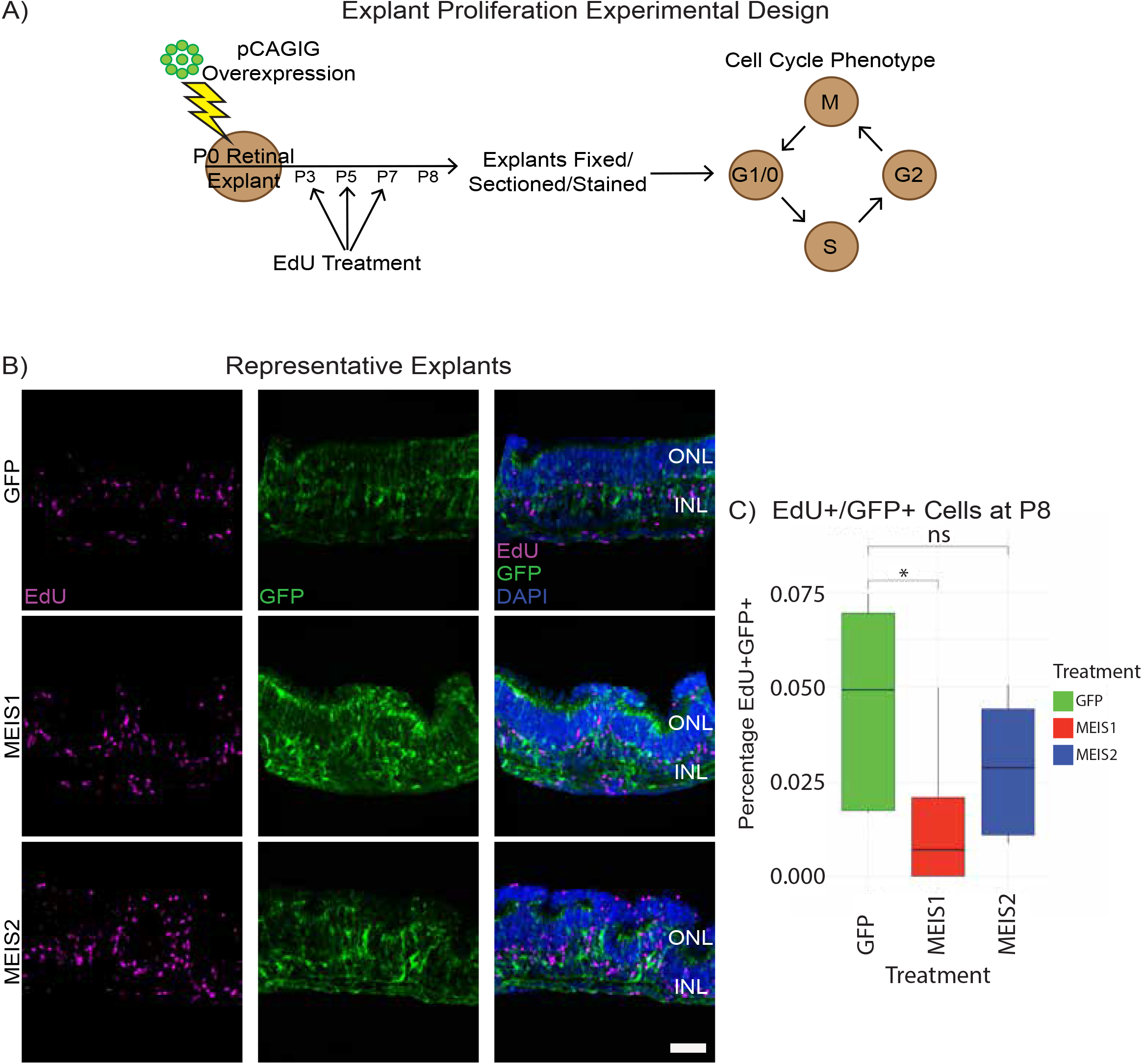
Impact of MEIS overexpression on retinal explant proliferation. A) Schematic showcasing the experimental design to analyze effects of *MEIS1* and/or *MEIS2* overexpression on RPC proliferation. B) Representative explants stained for EdU incorporation, GFP to represent electroporated cells, and DAPI for nuclei. C) Box plots to show the percentage of electroporated cells that are positive for EdU incorporation. P-values are indicated by * number. ns = not significant and * p< 0.05. Scale bar=50μm.

Finally, early-stage RPCs exhibit higher proliferation rates and symmetric self-renewing divisions (Turner, Snyder, and Cepko 1990). To examine whether MEIS overexpression affected RPC proliferation, we analyzed cell cycle phase markers (Fig. S12). At P2, primary RPCs were predominantly in G1, with a smaller G2 population. At P5, the proportion of G1 cells increased as expected. *MEIS1* increased G2-phase cells at P2 but reduced them at P5, whereas *MEIS2* had no effect. In neurogenic RPCs, all *MEIS* overexpression conditions increased the fraction of S/G2 transition cells at P2, with *MEIS2* maintaining this effect at P5. To directly assess proliferation, we performed EdU labeling at P3, P5, and P7, analyzing GFP+ EdU+ cells at P8 (Fig. 6a,b). *MEIS1* significantly reduced EdU incorporation, whereas *MEIS2* had no effect (Fig. 6c).

**Figure 6:**
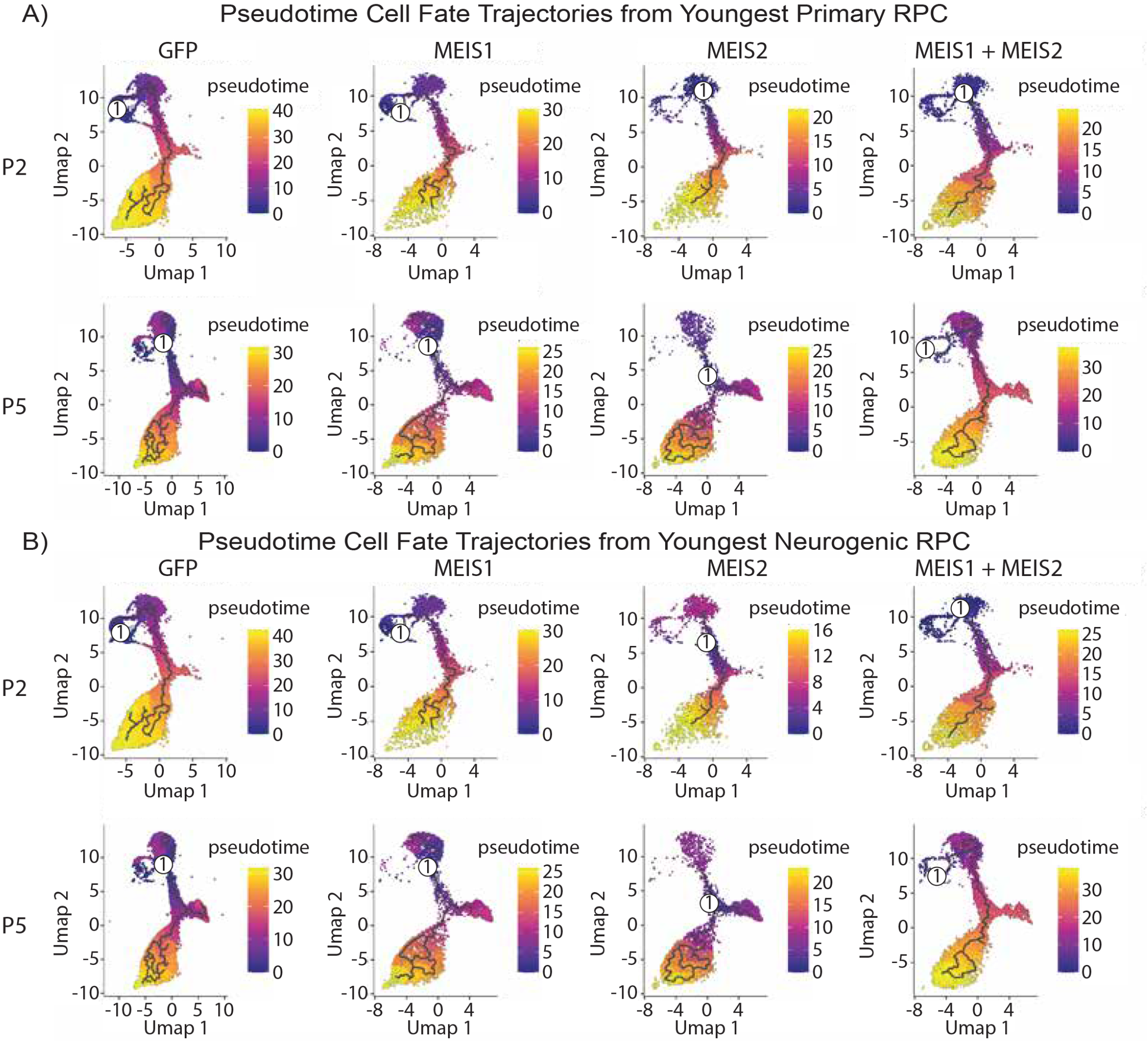
Monocle generated pseudotime trajectories for late born cell types present within the dataset. A) Using AUCell, a “youngest” primary RPC or B) neurogenic RPC was identified for each treatment dataset based on expression of known marker genes. This cell was used to generate maturation trajectories for each treatment dataset.

## Discussion

This study systematically examines the effects of ectopically expressing *Meis1* and *Meis2* in late-stage retinal progenitors, both individually and in combination. In mice, *Meis1* and *Meis2* are primarily expressed in early-stage retinal progenitor cells (RPCs), whereas in humans, *MEIS2* exhibits a distinct expression pattern, with increased levels in late-stage primary and neurogenic RPCs. Despite differences in expression dynamics, genes predicted to be repressed by *MEIS1/2* are consistently upregulated in late-stage RPCs in both species, whereas genes predicted to be activated by these factors show only modest early-stage enrichment. This discrepancy may reflect the divergent expression pattern of *MEIS2* between mouse and human.

Although *Meis1/2* have been implicated in promoting early-stage temporal identity, their overexpression in postnatal RPCs did not alter cell fate specification. However, significant changes in gene expression were observed, particularly in genes known to regulate retinal temporal patterning and neurogenesis. *MEIS1* selectively upregulated *Meis1* but had no effect on *Meis2*, whereas *MEIS2* activated *Meis2* but did not regulate *Meis1*. Both *MEIS1* and *MEIS2* robustly induced the early-stage marker *Hmga1*, but otherwise activated largely non-overlapping gene sets. For instance, *MEIS1* specifically activated the Notch effectors *Hes1* and *Hes5*, while *MEIS2* induced genes associated with both early (*Pbx1, Foxp4*) and late-stage (*Nfib*) temporal identity. Despite these differences, genes regulated by both factors were functionally enriched for neurogenesis, gliogenesis, apoptosis, and metabolic processes, such as glycolysis. The strong induction of *Hk1, Eno1b*, and *Eno1* following overexpression of *MEIS1* and/or *MEIS2* suggests a potential role for these factors in regulating metabolic states in early-stage RPCs.

While neither *MEIS1* nor *MEIS2* independently induced early-born cell types, they selectively modulated gene expression in ways that provide insight into their developmental functions. Both factors upregulated early-stage RPC-specific genes but failed to repress late-stage markers. Surprisingly, *MEIS2* even activated late-stage genes such as *Nfib*. In addition, *MEIS2* repressed Müller glia-specific genes while upregulating genes associated with cones, amacrine cells, and bipolar interneurons—cell types that endogenously express *Meis2* during differentiation. These findings suggest that *MEIS2* can activate lineage-specific transcriptional programs without fully driving cell fate commitment. Notably, *MEIS2*, but not *MEIS1*, accelerated the differentiation of postnatally generated neurons, further supporting its role in promoting neuronal maturation. Single-cell RNA sequencing suggested that *MEIS1* promotes primary RPC proliferation while *MEIS2* enhances neurogenic RPC proliferation. However, histological analysis revealed a reduction in EdU-labeled cells at P8, suggesting that overexpression of these factors does not sustain proliferation and may instead lead to cell cycle exit or cell loss.

Taken together, these findings demonstrate that *MEIS1* and *MEIS2* overexpression can activate a subset of early-stage RPC genes and promote expression of select early-born neuronal markers, yet they are insufficient to fully reprogram temporal identity or alter the fate of late-born retinal cell types. This is despite their known necessity for initiating neurogenesis and generating early-born cell types (Heine et al. 2008; Dupacova et al. 2021; Islam et al. 2013; Marcos et al. 2015). Several factors may account for these limitations. First, *Meis1* and *Meis2* function within a complex gene regulatory network governing early-stage temporal identity (Lyu et al. 2021). The absence of essential cofactors or intrinsic partners normally present in early-stage RPCs may constrain their ability to reprogram late-stage progenitors. Second, the fact that the *ex vivo* electroporation protocol used here excludes extraretinal cell types, as well as retinal pigment epithelium, may result in the lack of critical extrinsic signals required for efficient fate reprogramming. Third, both *MEIS1* and *MEIS2* upregulate proapoptotic factors, potentially leading to selective loss of cells that undergo more efficient temporal reprogramming.

This study highlights the challenges of using individual transcription factors to reprogram temporal identity in retinal progenitors. These findings have implications for regenerative strategies aimed at converting pluripotent stem cells or endogenous retinal glia into photoreceptors and other neuron types lost in blinding diseases. Understanding the broader regulatory networks governing temporal patterning may be key to overcoming these limitations and enhancing the success of cell reprogramming approaches.

## Data availability

All scRNA-Seq data are available in GEO as GSE295688.

## Acknowledgements

This work was supported by NIH grant R01EY036173 to S.B, and to F31EY033207 to P.L.

## Supplemental Figures

**Supplemental Figure 1: Heatmap showing expression of *Meis1/2*, and expression of genes predicted to be directly upregulated or downregulated by Meis factors in mouse retina**. A) Primary RPCs and B) neurogenic RPCs were analyzed from the dataset published in Clark et al 2019. Pheatmap expression plot is shown for *Meis1, Meis2*, and predicted targets of Meis1/2 that were identified in Lyu et al 2021.

**Supplemental Figure 2: *MEIS1/2* and expression of genes predicted to be directly upregulated or downregulated by Meis factors in human retina**. A) Primary RPCs cells and B) neurogenic RPCs were isolated from the dataset published in Zuo et al 2024. Violin expression plots for *MEIS1* and *MEIS2* and predicted targets of Meis1/2 that were identified in Lyu et al 2021.

**Supplemental Figure 3: *MEIS1/2* gene and predicted *MEIS1/2* target gene expression in human retinal progenitors**.

A) Primary retinal progenitor cells and were isolated from the dataset published in Zuo et al 2024. Pheatmap expression plot for MEIS1, MEIS2, and the components of the predicted upregulated and down regulated GRN from Lyu et al 2021 across developmental ages. B) Neurogenic retinal progenitor cells were isolated from the Zuo et al 2024 dataset. Pheatmap expression for the same genes as shown in A across the same developmental time points.

**Supplemental Figure 4: GFP expression in electroporated retinal explants**.

A) Representative fluorescent images of retinal explants 2 days and 5 days post electroporation with the pCAGIG overexpression construct for each treatment group. B) Representative FACS data for the GFP expression in electroporated explants with the percentage of GFP+ cell sorted for scRNA-analysis shown. C) Feature plot expression for GFP within the scRNA-seq expression. D) Violin plot for GFP expression for each cell type present in the scRNA-seq dataset. E) Violin plot for GFP expression for each treatment group present in the scRNA-seq dataset.

**Supplemental Figure 5: Uniform manifold approximation and projection (UMAP) images for each of the individual treatment groups from the scRNA-Seq dataset**.

This is generated from the dataset shown in Figure 2A.

**Supplemental Figure 6: Impacts of combined *MEIS1*+*MEIS2* overexpression on gene expression**. A) Volcano plot generated from differentially expressed genes between the P2 and P5 control treated samples and the P2 and P5 *MEIS1* + *MEIS2* overexpressing samples. B) Venn diagram showcasing the overlap in the genes expressed at a 0.5 Log2 Fold change between P2 and P5 *MEIS1, MEIS2*, and *MEIS1* + *MEIS2* overexpressing samples.

**Supplemental Figure 7: AUCell scoring for terminal cell fates in the primary RPC population *MEIS1/2* overexpression**. A) Violin scoring plot for retinal ganglion cell marker genes. B) Violin scoring plot for shared amacrine and horizontal cell marker genes. C) Violin scoring plot for cone photoreceptor marker genes. D) Violin scoring plot for rod photoreceptor marker genes. E) Violin scoring plot for bipolar cell marker genes. F) Violin scoring plot for Muller glia marker genes. P-values are indicated by * number. ns = not significant, * p< 0.05, ** p< 0.01, *** p< 0.001, and **** p< 0.0001.

**Supplemental Figure 8: AUCell scoring for terminal cell fates in neurogenic RPCs population *MEIS1/2* overexpression**. A) Violin scoring plot for retinal ganglion cell marker genes. B) Violin scoring plot for shared amacrine and horizontal cell marker genes. C) Violin scoring plot for cone photoreceptor marker genes. D) Violin scoring plot for rod photoreceptor marker genes. E) Violin scoring plot for bipolar cell marker genes. F) Violin scoring plot for Muller glia marker genes. P-values are indicated by * number. ns = not significant, * p< 0.05, ** p< 0.01, *** p< 0.001, and **** p< 0.0001.

**Supplemental Figure 9: Heatmap showing differential expression of *Meis1/2*, and genes predicted to be either upregulated or downregulated by Meis factors in primary and neurogenic RPCs following *MEIS1/2* overexpression**.

**Supplemental Figure 10: Primary RPC expression of early and late born cell type markers following *MEIS1/2* overexpression**. A) Pheatmap for marker genes associated with early-born retinal cell types and B) late-born cell types in P2 and P5 primary RPCs. Abbreviations: epRPCs: early primary retinal progenitor cells, enRPCs: early neurogenic RPCs, RGCs: retinal ganglion cells, ACs: amacrine cells, HCs: horizontal cells, lpRPCs: late primary RPCs, lnRPCs: late neurogenics RPCs, BCs: bipolar cells, MG: Muller glia.

**Supplemental Figure 11: Neurogenic RPC expression of early and late born cell type markers following *MEIS1/2* overexpression**. A) Pheatmap for marker genes associated with early-born cell types and B) late-born cell types in P2 and P5 neurogenic RPCs. Abbreviations: epRPCs: early primary retinal progenitor cells, enRPCs: early neurogenic RPCs, RGCs: retinal ganglion cells, ACs: amacrine cells, HCs: horizontal cells, lpRPCs: late primary RPCs, lnRPCs: late neurogenics RPCs, BCs: bipolar cells, MG: Muller glia.

**Supplemental Figure 12: Bioinformatically calculated cell stage densities for primary and neurogenic RPCs at P2 and P5**. A) Density line plots for P2 and P5 primary RPCs and B) neurogenic RPCs for all cell cycle stages.

**Supplemental Figure 13: Monocle generated pseudotime trajectories for late born cell types within the dataset**. A) Using AUCell, the “oldest” primary RPC and B) neurogenic RPC was identified using was identified for each treatment dataset based on expression of known marker genes. This cell was used to generate maturation trajectories for each dataset.

## Supplemental Tables

**Supplemental Table 1**: List of cell type-specific markers used for analysis, and changes in aggregate expression of these markers by treatment condition.

**Supplemental Table 2**: List of genes differentially expressed in GFP-positive cells overexpressing *MEIS1, MEIS2*, or *MEIS1+MEIS2* relative to GFP-positive cells electroporated with the pCAGIG control plasmid.

## Methods

### Mice

The use of animals for these studies was conducted using protocols approved by the Johns Hopkins Animal Care and Use Committee, in compliance with ARRIVE guidelines, and were performed in accordance with relevant guidelines and regulations. Timed pregnant mice were ordered from Charles River Laboratories for the retinal explant electroporation experiments ending in scRNA-seq or histological analysis.

### Immunohistochemistry and imaging

P8 retinal explants were fixed in 4% paraformaldehyde (Electron Microscopy Sciences, no. 15710) for 2 hours at 4°C, then carefully detached from the membrane in 1× PBS and incubated in 30% sucrose overnight at 4°C. The explants were then embedded in frozen section medium (VWR, no. 95057-838), cryosectioned at 16-μm thickness, and stored at −20°C. Sections were dried for 20 min in a 37°C incubator and washed three times for 5 min with 0.2% Triton X-100 in PBS (PBST). EdU labeling was performed using a Click-iT EdU kit (Thermo Fisher Scientific, no. C10340) following the manufacturer’s instructions. Sections were then incubated in blocking buffer (10% horse serum (Thermo Fisher Scientific, no. 26050070), 0.2% Triton X-100 in 1× PBS) for 1 hour at room temperature (RT) and subsequently incubated with primary antibodies in blocking buffer overnight at 4°C.

Sections were washed three times for 5 min in PBST to remove excess primary antibodies and were then incubated in blocking buffer with secondary antibodies for 1 hour at RT. The sections were washed three times for 5 min in PBST, then once for 5 minutes in PBS. Sections were then mounted with Fluoromount-G™ with DAPI (Thermo Fisher Scientific, no. 00-4959-52) under coverslips (VWR, no. 48404-453), air-dried, and stored at 4°C. Fluorescent images were captured using a Zeiss LSM 700 confocal microscope.

### Single-cell RNA-sequencing (scRNA-seq) library preparation

ScRNA-seq were prepared on GFP+ cells isolated from dissociated retinal explants using the 10X Genomics Chromium Single Cell 3’ Reagents Kit v3.1 (10X Genomics, Pleasanton, CA). Libraries were constructed according to manufacturer’s instructions and were sequenced using Illumina NextSeq. Sequencing data were processed through the Cell Ranger 7.0.1 pipeline (10X Genomics) using default parameters.

### scRNA-seq data analysis

ScRNA-Seq data were pre-processed using Cell Ranger v7.1.0 using custom mouse genome mm10 with an added GFP sequence. The cell-by-genes count matrices were further analyzed using Seurat V5.1.0 (Hao et al. 2024).Cells with RNA counts less than 500 or greater than 50,000, and number of genes less than 200 or greater than 6000 were filtered out as low quality cells. Additionally, cells with a mitochondrial fraction of greater than 25% were removed. Lastly, cells with a GFP expression level of less than 1 were removed. For data visualization, UMAP was generated using the first 22 dimensions. Primary RPC and neurogenic RPC cell clusters were identified based on expression of published marker genes and subsetted for further analysis. Differential gene expression between the control group and *MEIS1, MEIS2*, or *MEIS1* + *MEIS2* was determined through Seurat’s built-in FindMarkers function.

### Dataset analysis R packages

AUCell was used for the aggregating scoring plots in Figures 1 and 4, and supplemental figures 2, 7, and 8 (Aibar et al. 2017). Pheatmap was used to generate the heatmaps in Supplemental Figures 1, 3, 9, 10, and 11 (Kolde 2012). ggplot2 was used to generate the Volcano Plots shown in Figure 3 and Supplemental Figure 6 (Wickham 2009). ggplot2 was used to generate the GO term enrichment plots shown in Figure 3 (Wickham 2009). The Venn Diagram plots generated for Figure 3 and Supplemental Figure 6 created with ggvenn (“[No Title],” n.d.). ggplot2 was used to generate the box plot for the EdU incorporation in Figure 5 (Wickham 2009). Tricycle was used to generate the cell cycle stage density plots in Supplemental Figure 12 (Zheng et al. 2022). Monocle3 was used to generate the cell fate trajectory plots shown in Figure 6 and Supplemental Figure 13 (Cao et al. 2019).

