## Supplemental Figures 1-13 for "Overexpression of Meis factors in late-stage retinal progenitors yields complex effects on temporal patterning and neurogenesis"

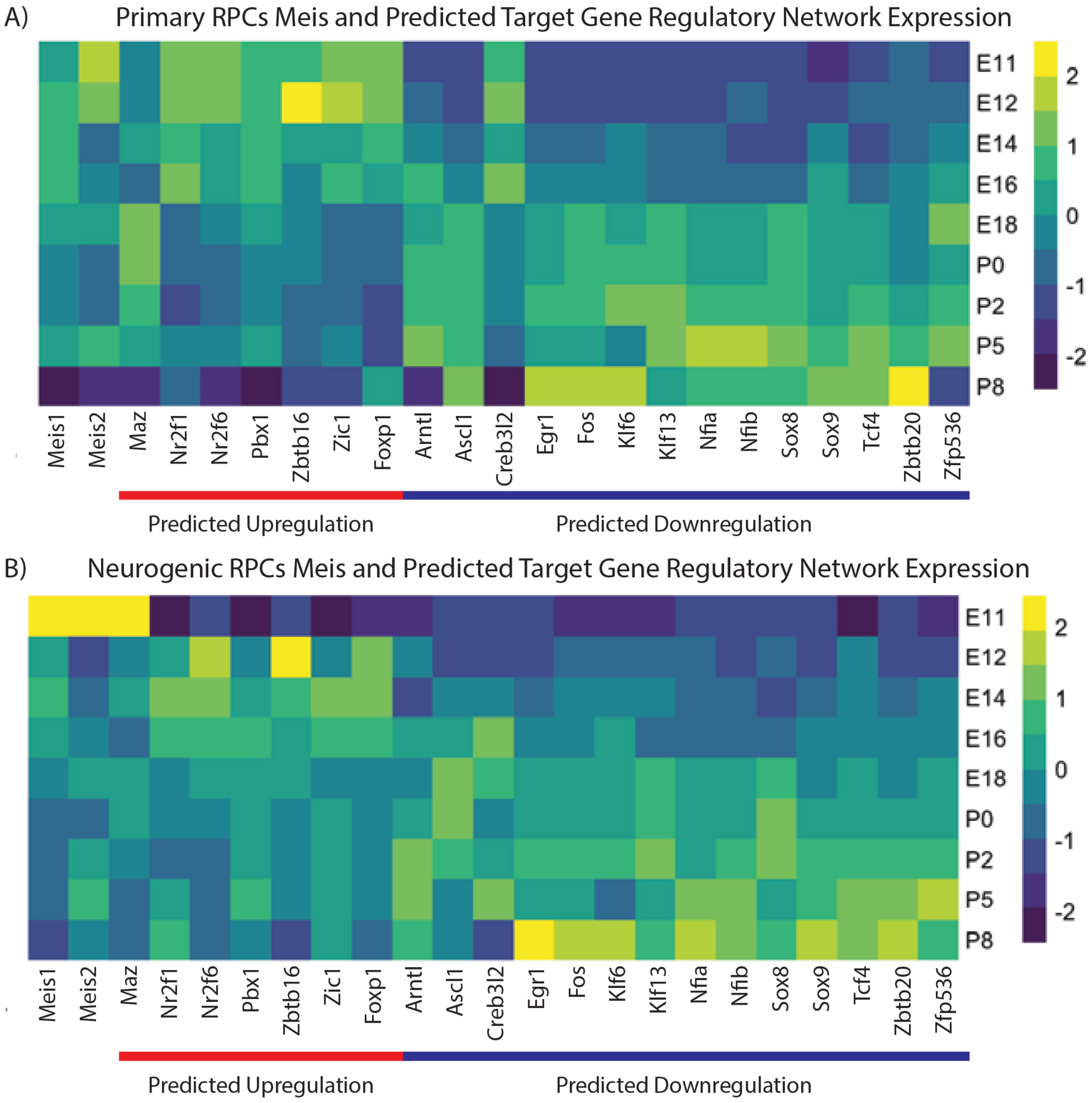


Leavey, et al. Fig. S1

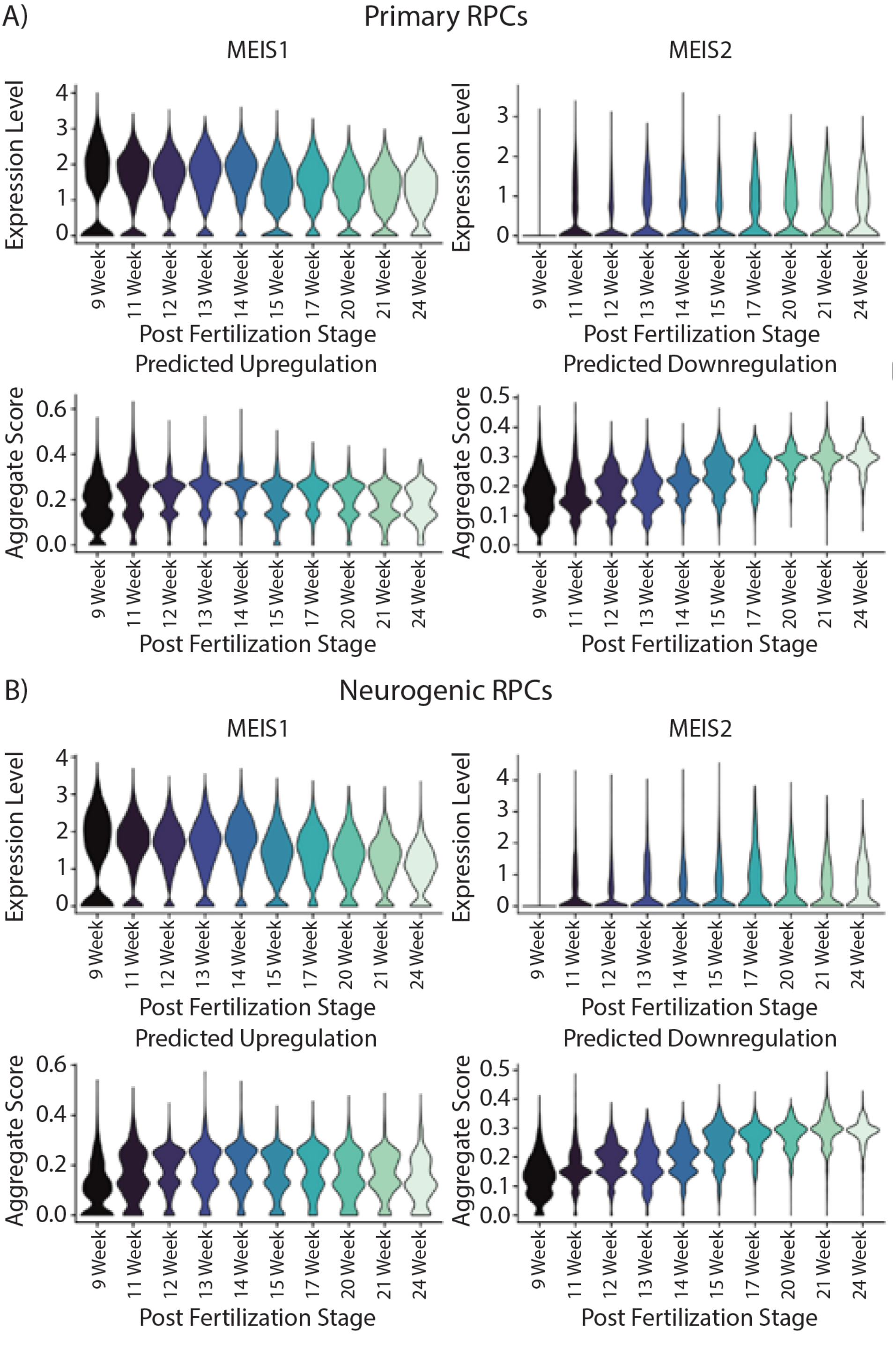


Leavey, et al. Fig. S2

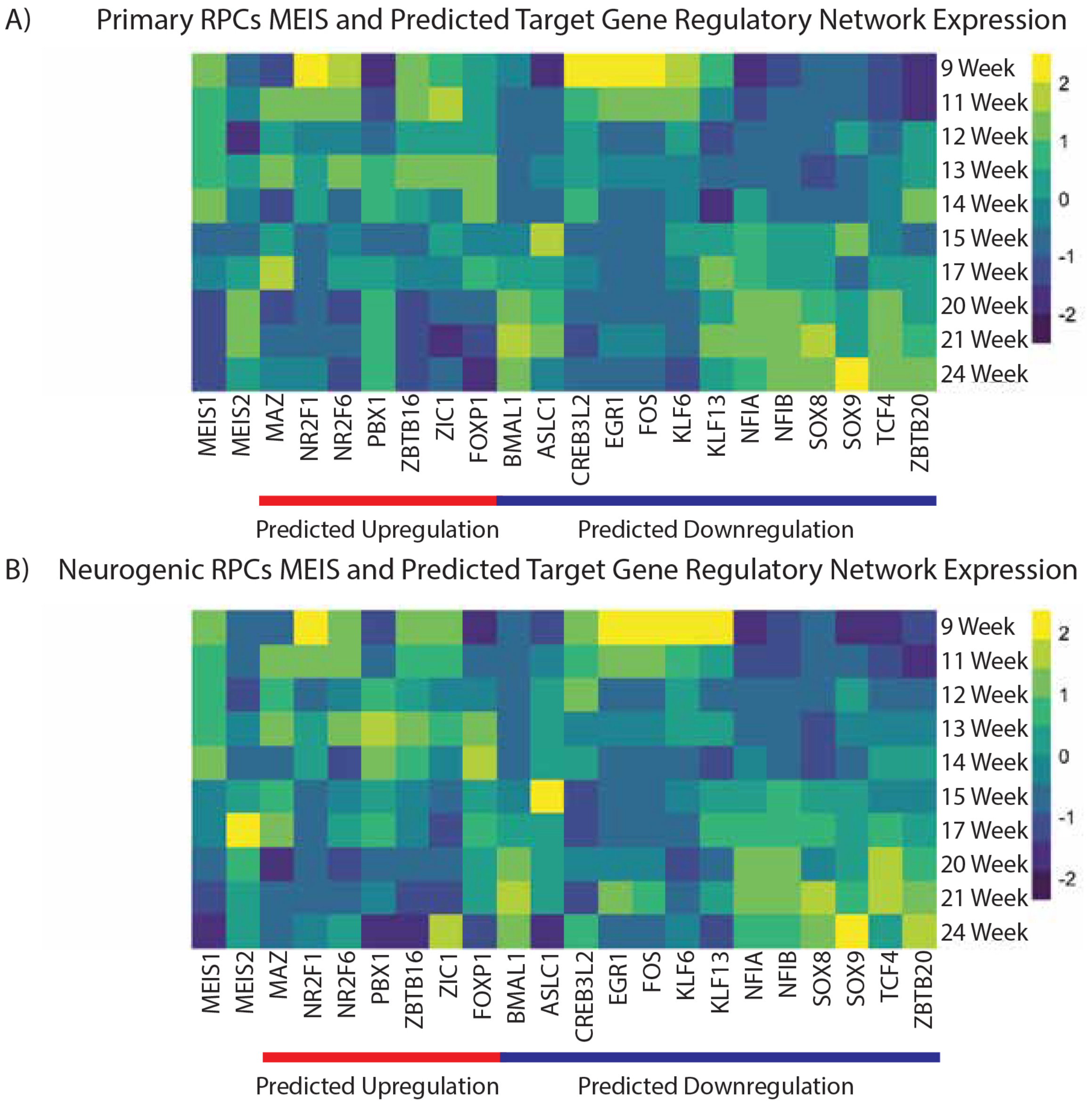


Leavey, et al. Fig. S3

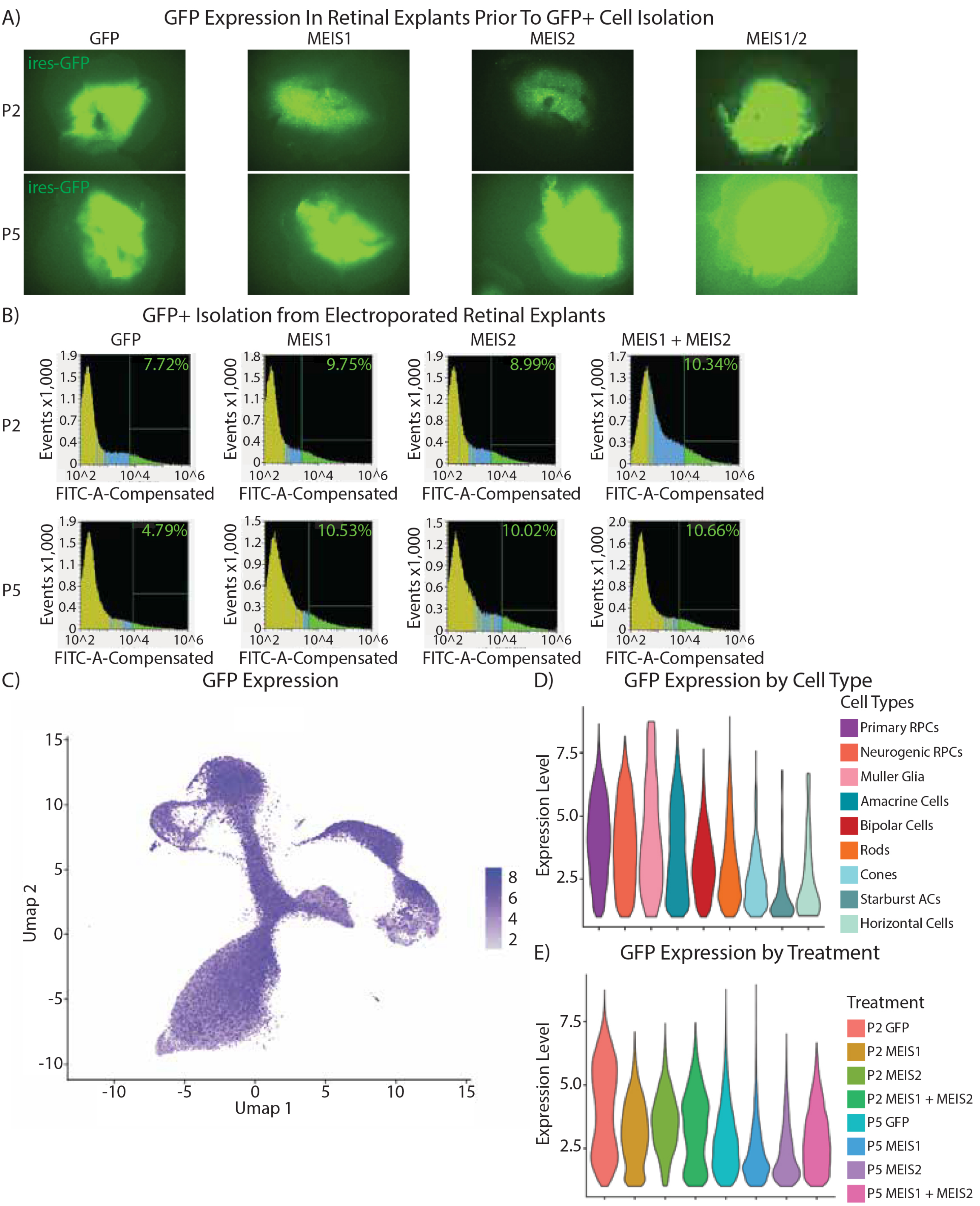


Leavey, et al. Fig. S4

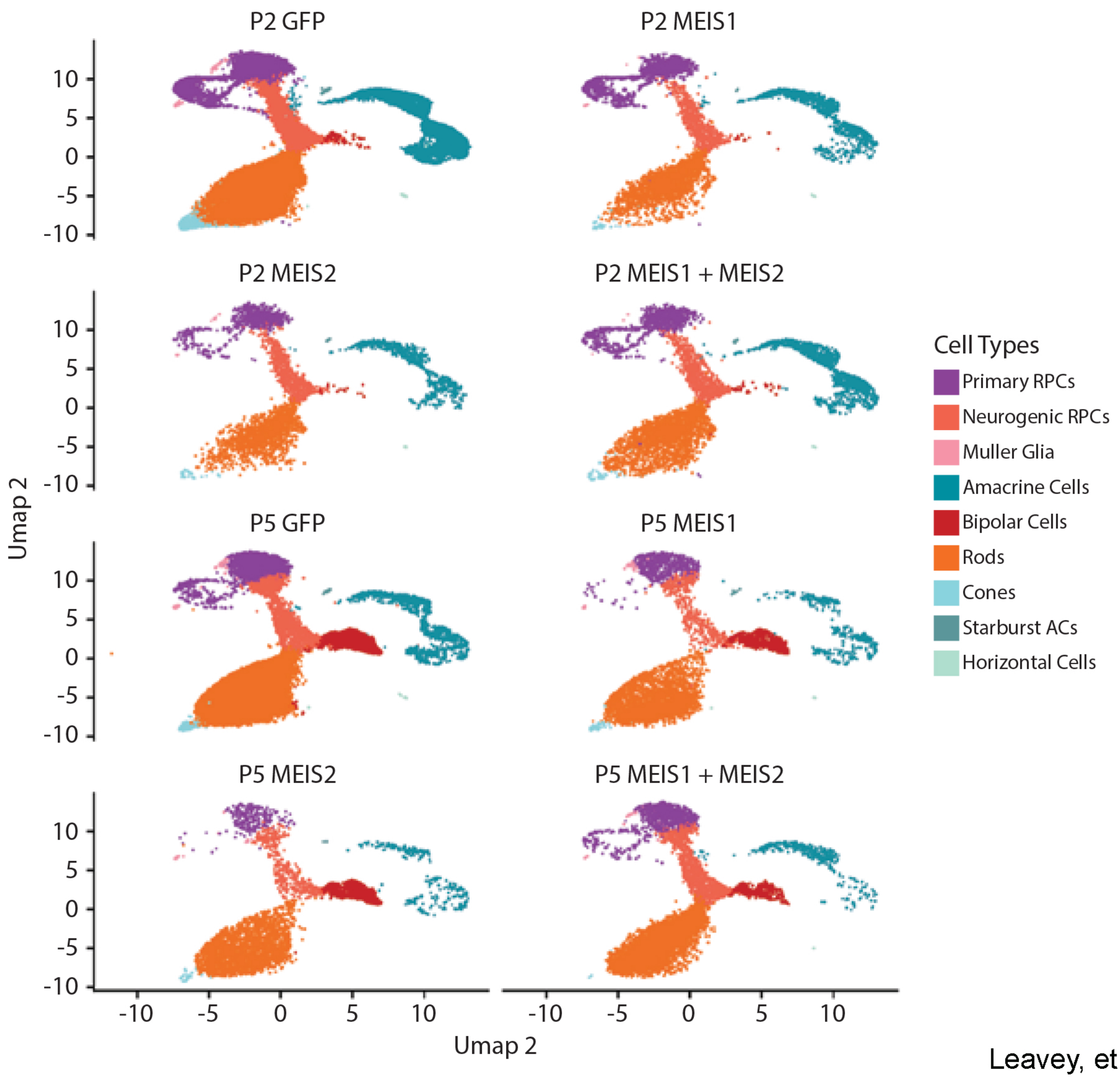


Leavey, et al. Fig. S5

A) MEIS1 + MEIS2 Enriched Gene Expression

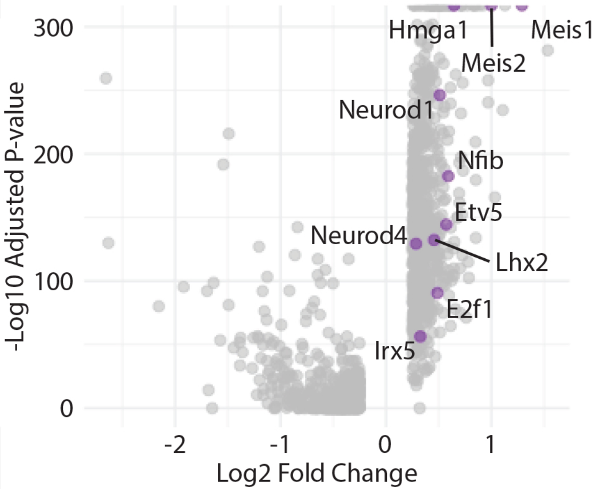


B) Differentially Expressed Gene Overlap for MEIS1 MEIS2 and MEIS1 + MEIS2

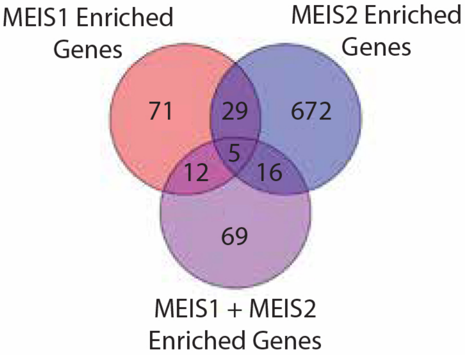


Leavey, et al. Fig. S6

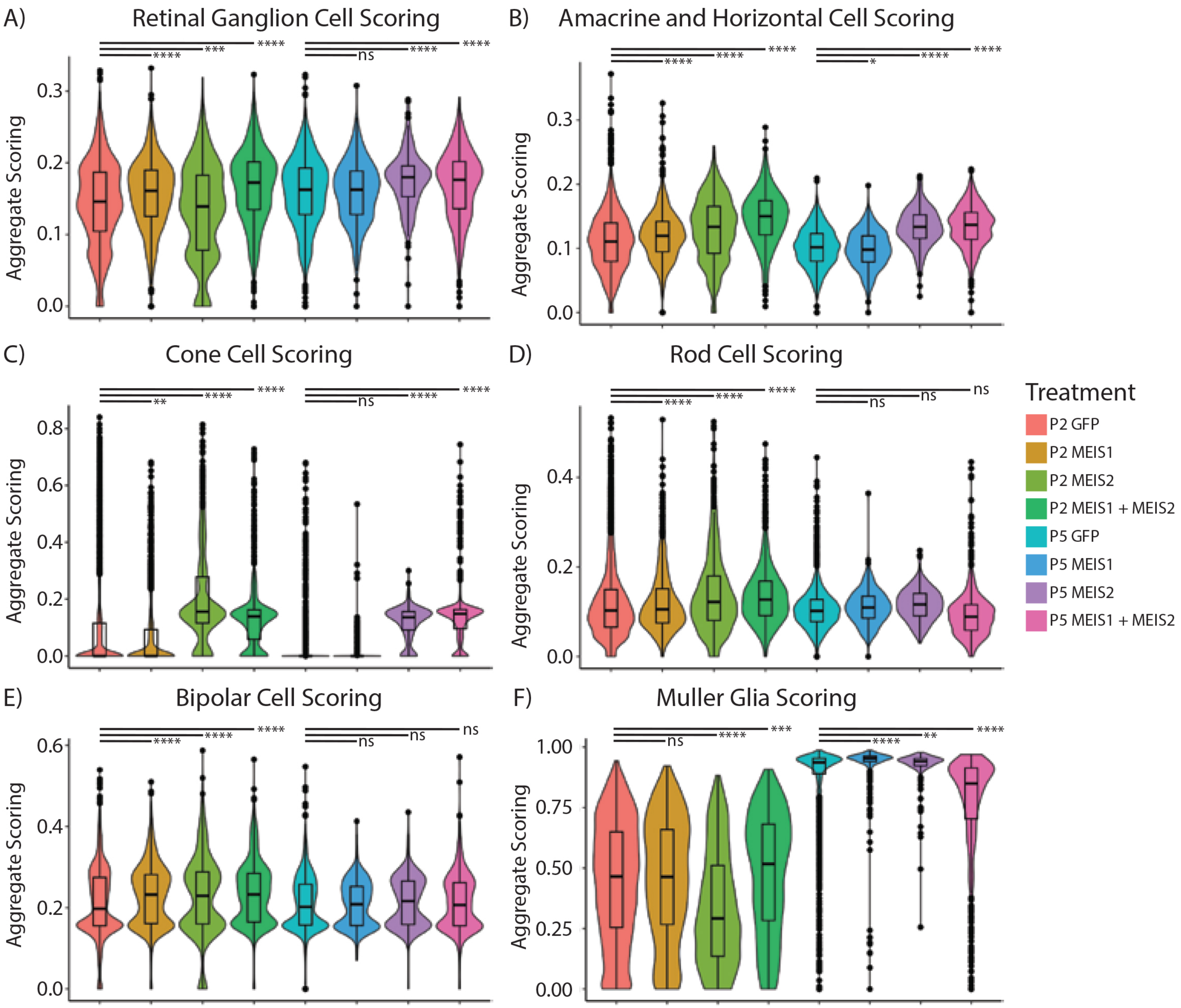


Leavey, et al. Fig. S7

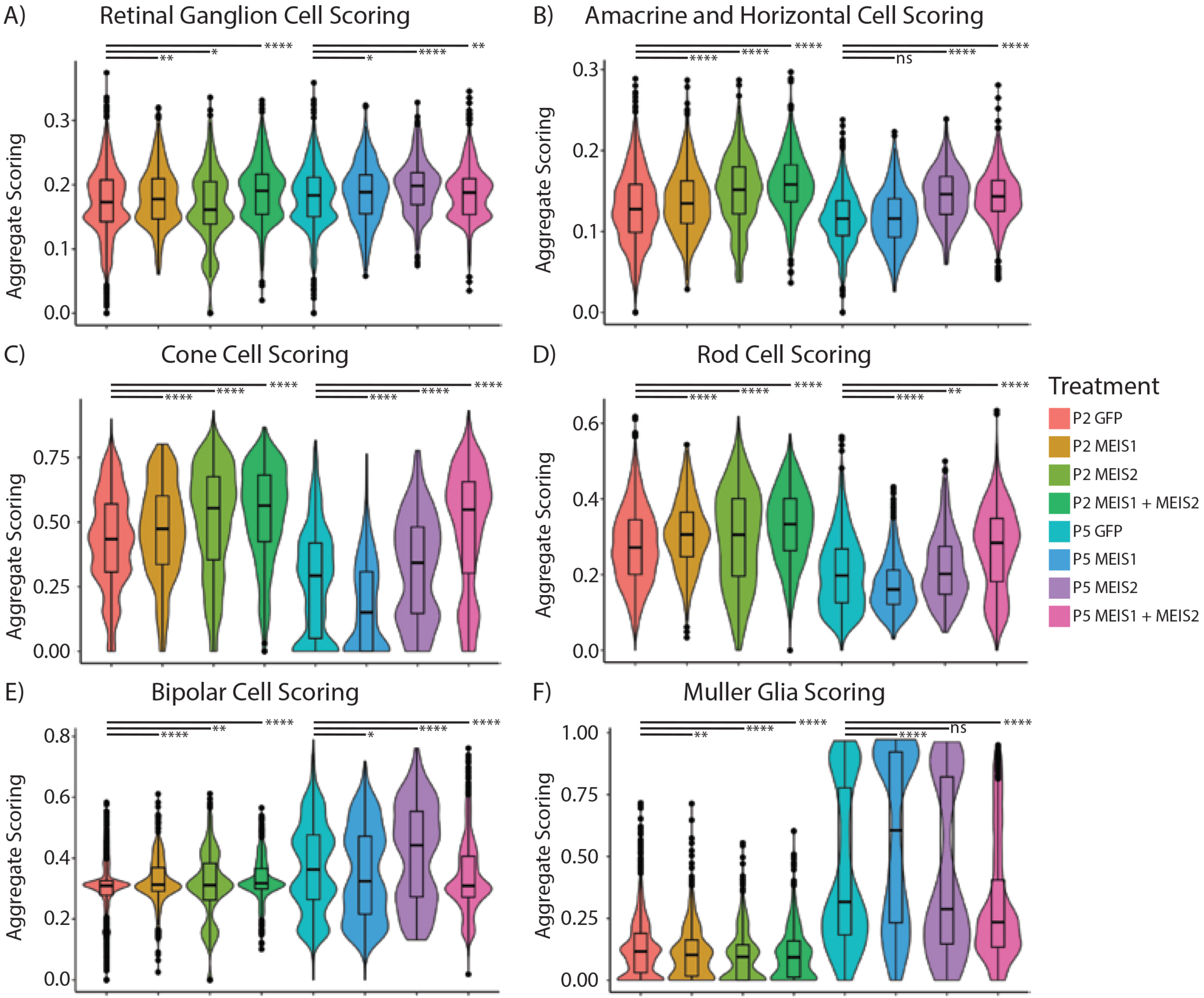


Leavey, et al. Fig. S8

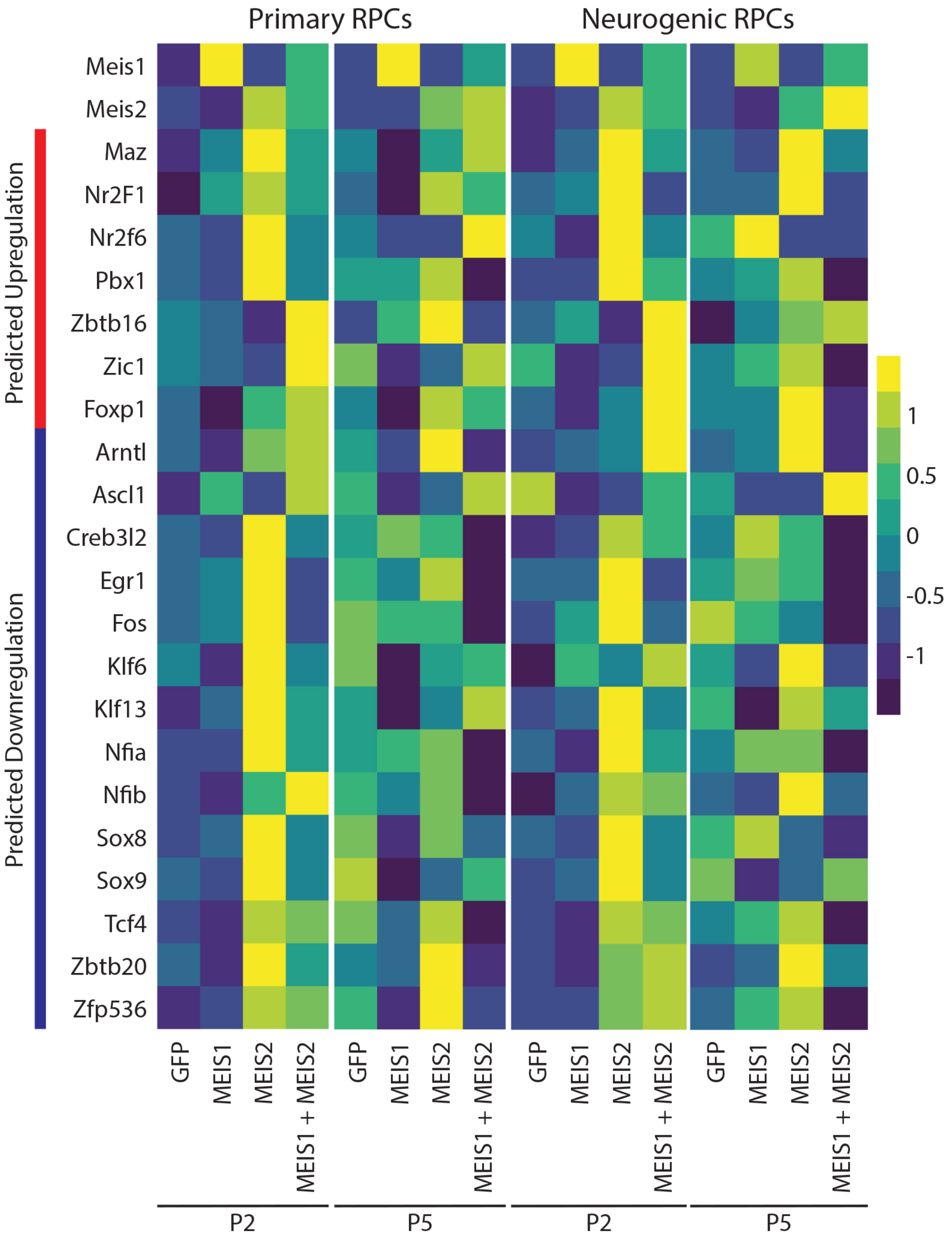


Leavey, et al. Fig. S9

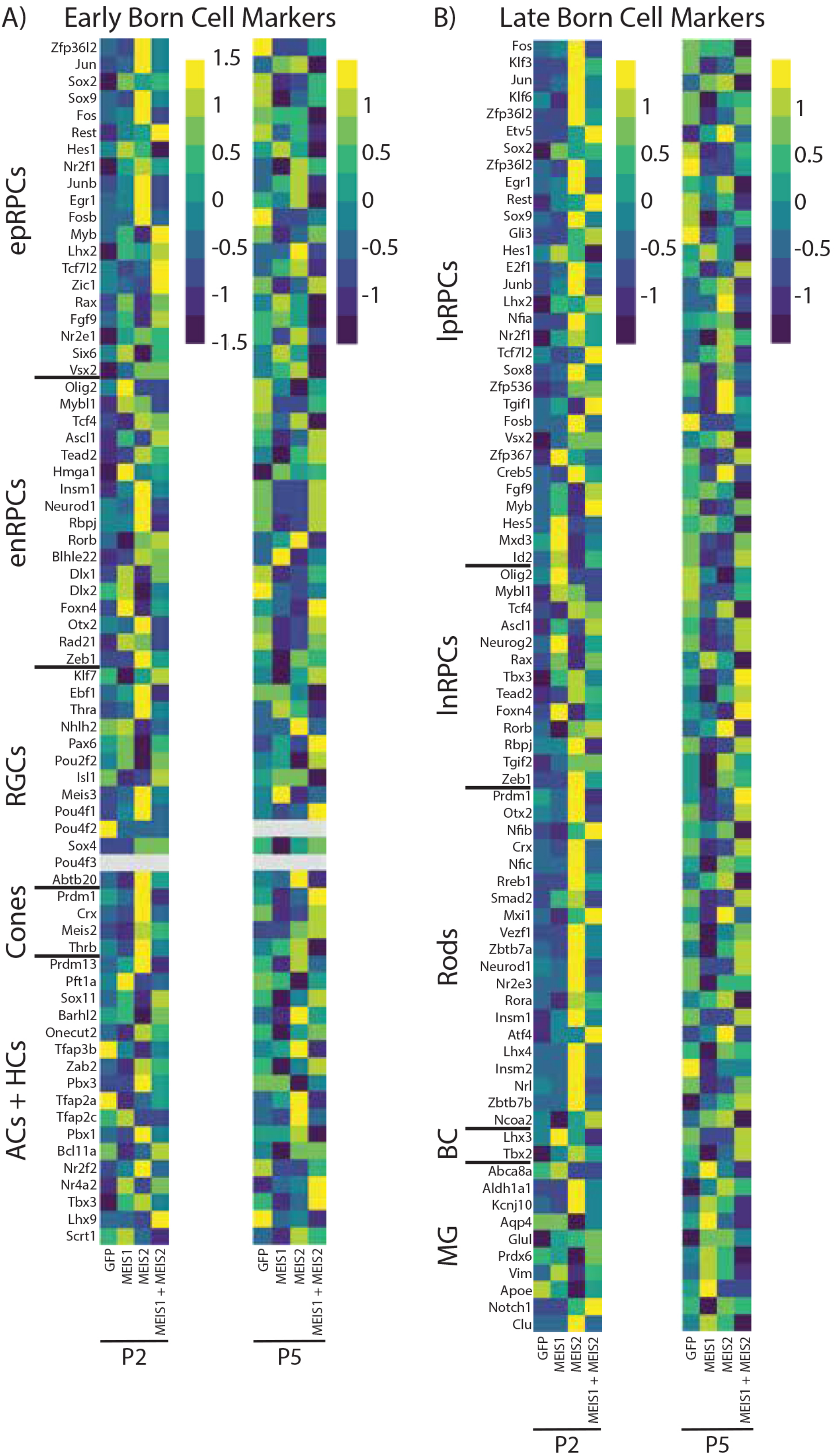


Leavey, et al. Fig. S10

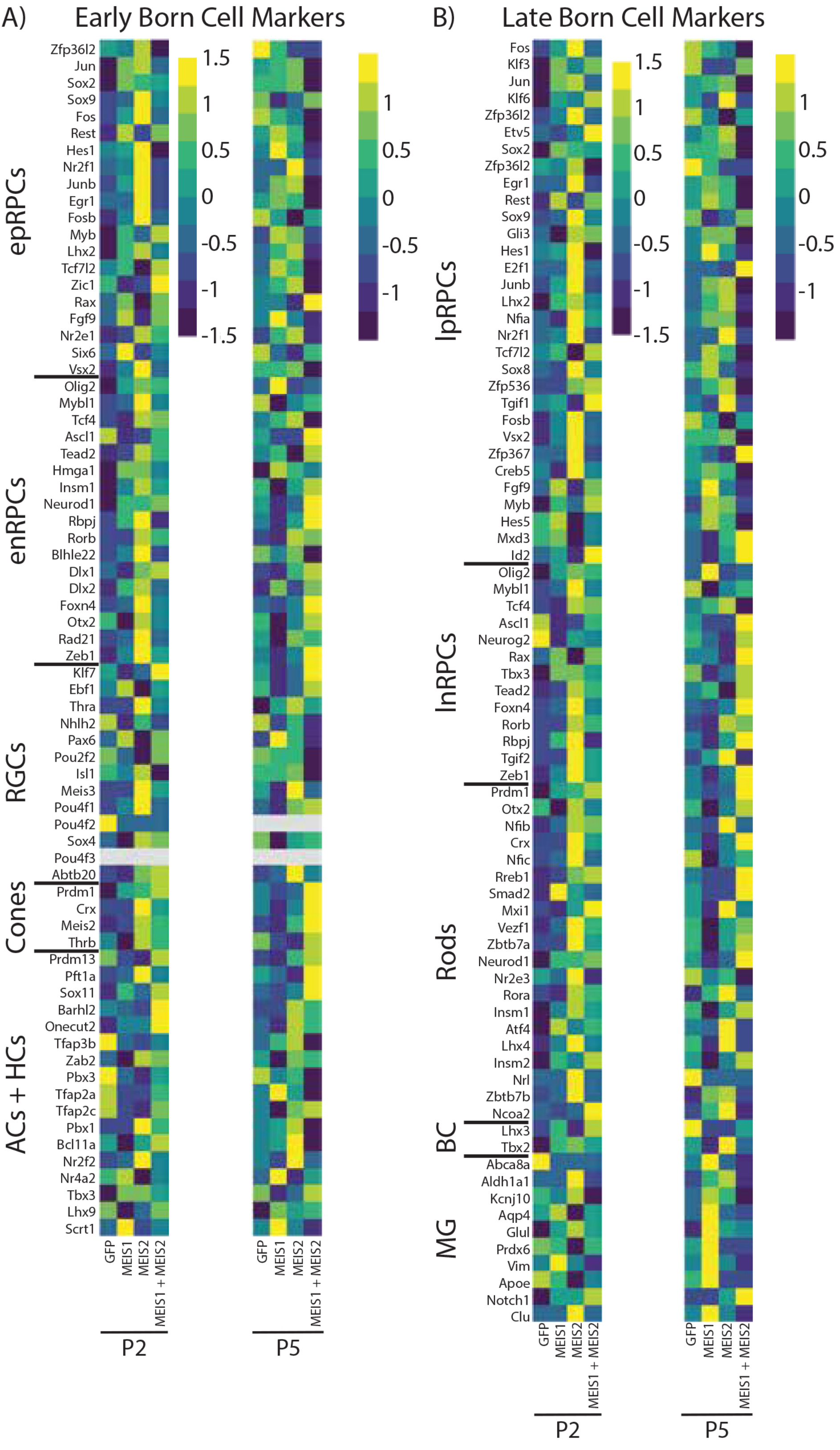


Leavey, et al. Fig. S11

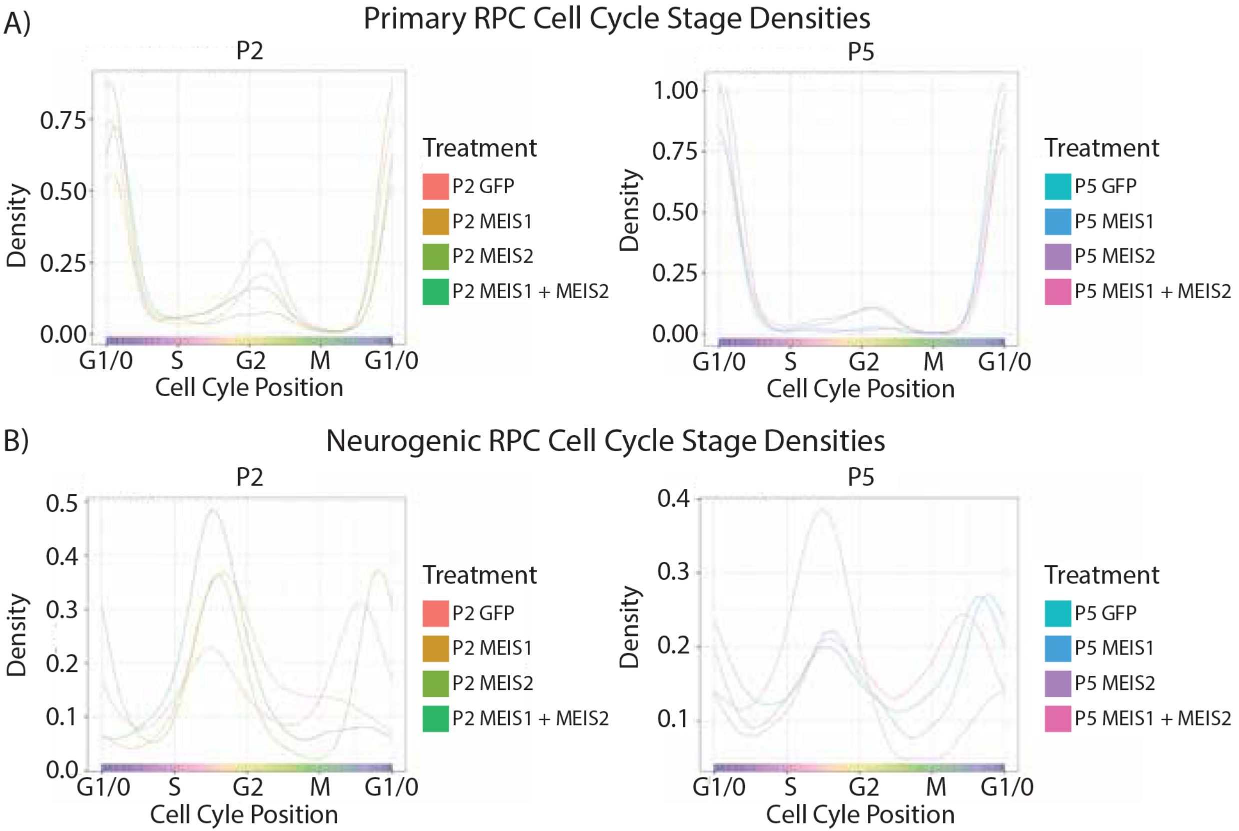


Leavey, et al. Figure S12

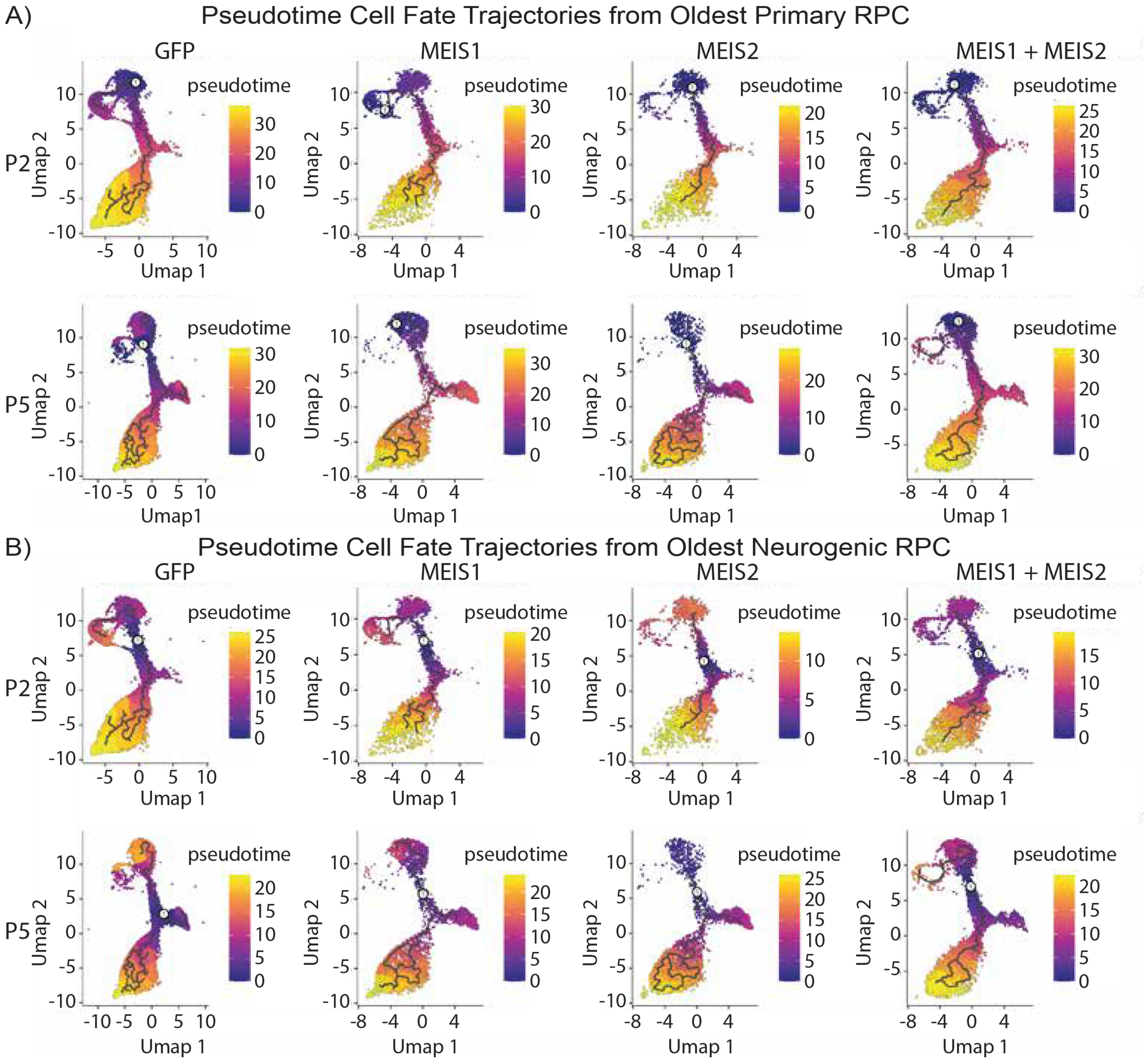


Leavey, et al. Figure S13
